# GSPlot: Visualizing gene set similarity in enrichment results

**DOI:** 10.64898/2026.06.11.731691

**Authors:** Tien Cat Le, Cristian Gutierrez Espinoza, Jason Courtois, Özgün Babur

## Abstract

**Summary:** Interpretation of gene set enrichment analyses is challenging when the results contain many sets. Similarities in those sets can be used for complexity management. We present GSPlot, a web-based application for visualizing enrichment results of large gene set collections. GSPlot displays gene sets in a two-dimensional space where distances reflect the similarity of their enriched content. This view brings a higher-level structure to the enrichment results and simplifies their interpretation and communication.

**Availability and implementation:** GSPlot is available at https://gsplot.cs.umb.edu/. Source code is available at https://github.com/PathwayAndDataAnalysis/gsplot.

## 1. Introduction

To interpret comparative high-throughput molecular profiles, biologists often use enrichment analyses over predefined gene sets (such as GSEA (Subramanian et al., 2005; Franz et al., 2025)), which connect molecular changes to biological processes, cellular pathways, and other phenomena that are represented by a set of genes. Gene set enrichment results are traditionally presented as a list where the names and p-values of the enriched gene sets are displayed. The list view is not easy to interpret if the result volume is high. In such cases, the results often contain highly overlapping gene sets, not necessarily having similar names. Gene set similarities provide an opportunity to organize their presentation into more comprehensible structures.

Gene set similarity was previously used to display enrichment results as a network of gene sets (Merico et al., 2010) where similar sets are connected with an edge. Here, we present GSPlot, an alternative approach to organizing enrichment results. GSPlot uses context-specific set similarity measures to plot the gene sets in 2D space, where the gene sets naturally form clusters. AI-assisted annotation of these clusters provides a high-level view of the enrichment results. This serves as a summary and facilitates the interpretation.

## 2. Methods

GPlot was implemented in Python as a web service using the Django and Plotly frameworks. The sections below describe implementation details and user options.

### 2.1 Gene set resource

Users can select a gene set collection in the Molecular Signatures Database (MSigDB) (Liberzon et al., 2011) or upload their custom collection.

### 2.2 Enrichment methods

We support three modes of enrichment testing, which correspond to three different gene input formats: ranked genes, thresholded genes, and scored genes. In all modes, prior to the analysis, we prune both the gene input and the gene sets to their intersection. This means filtering out the genes in the input that are not members of any gene set, and also filtering out the members of gene sets that are not present in the gene input. Pruning is a measure against a possible gene annotation bias in the analysis.

#### 2.2.1 Ranked genes

Users provide a ranked list of gene symbols that can be either unipolar (from most significant to least significant) or bipolar (from most up-regulated to insignificant, then to most down-regulated). The gene set enrichment direction and significance are determined through a mean rank test. Let *R*_*i*_ be the integer rank of the gene *i*. Its normalized rank *r*_*i*_ = (*R*_*i*_ − 0.5)*/n* where *n* is the number of genes. The normalization scales ranks in the range (0, 1). The mean rank of gene set *S*, calculated as *m*_*S*_ = (Σ _*i∈S*_ *r*_*i*_)*/k*, is the enrichment indicator where *k* = |*S*|. *m*_*S*_ *<* 0.5 means positive enrichment and *m*_*S*_ *>* 0.5 means negative enrichment. To assess its significance, we use the standard deviation of the mean rank 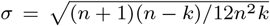 , assume the null distribution is a Gaussian around 0.5, convert the mean rank into a z-value *z* = (*m*_*S*_ − 0.5)*/σ*, and use the cumulative distribution function (Φ) of *N* (0.5, *σ*) to estimate the p-value. Φ(*z*) is one-tailed p-value for detecting positive enrichment, Φ(−*z*) is one-tailed p-value for detecting negative enrichment, and 2Φ(−|*z*|) is the 2-tailed p-value for detecting both types of enrichment together.

#### 2.2.2. Thresholded genes

Users provide two gene lists: genes with a significant change and genes without a significant change. We generate a contingency table for each gene set that compares significant and insignificant genes inside and outside of the gene set. Consequently, we use a one-tailed Fisher’s exact test to assess whether each gene set is enriched among the significant genes.

#### 2.2.3. Scored genes

Users provide genes and their scores in a two-column text file. We sort the genes by descending scores and run GSEApy (Fang et al., 2023) prerank to detect enriched gene sets. Positive, negative, and two-sided enrichment modes are supported.

In all cases, we use the Benjamini-Hochberg method to control the false discovery rate.

### 2.3. Gene set distance measures

We calculate pairwise distances between enriched gene sets based on their member genes using Jaccard distance (Eq. 1) or overlap coefficient distance (Eq. 2).

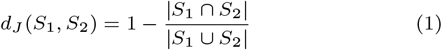

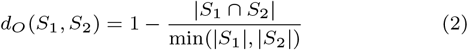

GSPlot also supports weighted versions of both distance measures (Eqs. 3-4) where *w*_*i*_ is the non-negative weight of gene *i*.

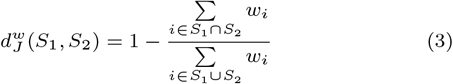

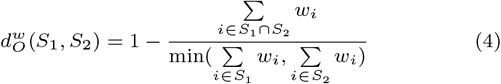

We use weighted distances to emphasize the genes that contributed to the enrichment. Gene weights depend on the input mode and enrichment type. In the case of ranked genes and scored genes, normalized ranks (*r*_*i*_) are used as gene weights for comparing negatively enriched sets, and 1 − *r*_*i*_ is used for comparing positively enriched sets. In the case of thresholded genes, significant gene weights are set to 1, and insignificant gene weights are set to 0. If two compared gene sets are enriched in opposite directions, then an unweighted distance is used because genes would have different importance in the two sets.

The resulting pairwise distances are then rescaled as *d*^*′*^ = *d/*(1−*d*), with an upper cap of 100 to avoid infinite values when two gene sets have no overlap. This rescaling increases the separation among weakly overlapping gene sets. The rescaled pairwise distances are input to the two-dimensional layout algorithms.

### 2.4. 2D layout of gene sets

GSPlot uses the distance matrix of the gene sets to generate a two-dimensional layout of the enriched gene sets. Users can choose among UMAP (Becht et al., 2019), t-SNE (Van der Maaten and Hinton, 2008), and Isomap (Tenenbaum et al., 2000) algorithms for the projection. In the resulting layout, each point represents an enriched gene set, and gene sets with smaller pairwise distances are placed closer.

### 2.5. Cluster detection

GSPlot supports cluster detection on the two-dimensional gene set map using HDBSCAN (Malzer and Baum, 2020) or OPTICS (Ankerst et al., 1999). Users can select one of the two algorithms and adjust its main parameters in the interface. The resulting cluster assignments are used for cluster coloring and cluster label generation.

### 2.6. Generation of cluster labels

For each detected cluster, GSPlot builds a compact cluster summary from the enriched gene sets assigned to that cluster. The summary includes the size of the cluster, the names of representative gene sets selected by the p-value, and the frequently occurring genes from the gene sets in the cluster. These summaries are sent to Gemini 2.5 Flash to generate a label for each cluster.

## 3. Results

We built GSPlot for performing and visualizing gene set enrichment analysis on large gene set collections. The method is especially useful when the results contain a large number of sets, such as tens or hundreds. GSPlot displays those sets in a 2D space where similar gene sets form clusters, and the clusters are annotated with summary labels. Clustering and the subsequent annotation provide an accessible entry point for interpreting the results.

To start an analysis, users need to select or upload a gene set collection and provide the gene input in one of the following forms: ranked genes, thresholded genes, or scored genes (Fig. 1A). Users also specify which enrichment direction they are interested in: positive, negative, or both. GSPlot uses an appropriate statistical method (see Methods) and identifies enriched gene sets. The resulting gene sets are then displayed in 2D space using one of the following 2D projection methods: t-SNE, UMAP, or Isomap. The selected projection algorithm decides the 2D locations of sets based on the set similarities. To assess pairwise set similarity, GSPlot uses Jaccard similarity or the overlap coefficient, either in plain or weighted modes (see Methods). In the weighted mode, a common gene with a higher contribution to enrichment brings a higher similarity to the gene sets, making the similarity measure enrichment-specific.

**Figure 1.**
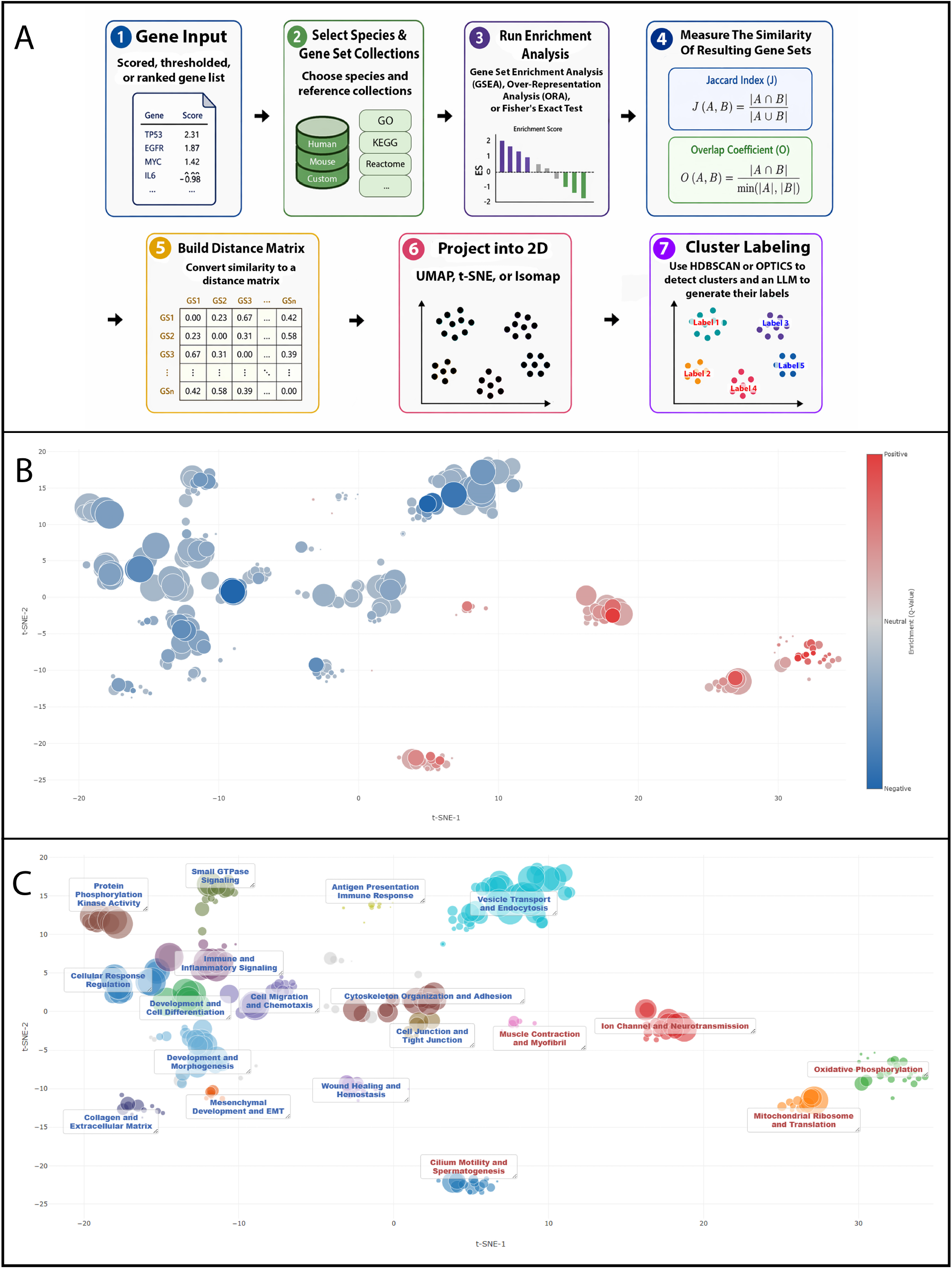
GSPlot workflow and visualization outputs. (A) Analysis workflow. (B) Significance-based view of the enrichment result. (C) Cluster view with labels colored by dominant enrichment direction.

In the 2D view, each point represents an enriched gene set, and the set area is correlated with the set size. Users can inspect the result in different visualization modes. In the significance-based view (Fig. 1B), point color indicates the enrichment direction, and its intensity is correlated with the strength of the enrichment. In the cluster view (Fig. 1C), colors indicate cluster membership and clusters are labeled. GSPlot uses HDBSCAN (Malzer and Baum, 2020) or OPTICS (Ankerst et al., 1999) algorithms for cluster detection, and uses the Gemini API to label the clusters. The cluster label color indicates the dominant enrichment direction within the cluster. Users are also provided with tools to further inspect the resulting gene sets. Clicking on a gene set displays the details of the enrichment and the member genes under the result plot. When multiple sets are selected, the common members are displayed in bold.

To demonstrate its utility, we reanalyzed data from a recent publication where aging cardiac cells are compared to cells from younger donors (Gao et al., 2026). In the original article, cell-type-specific differential RNA expressions are obtained and enrichments are calculated on the Hallmark (Liberzon et al., 2015) gene sets. The Hallmark gene set collection contains 50 curated gene sets with low overlaps; therefore, they are easy to interpret. However, this simplification comes with the cost of potential missing processes in the results. Comparing aging and young cardiomyocytes, the authors identify 6 sets (1 positive, 5 negative, in their Fig. S13d) and mention 2 of them in the Results section: positive enrichment in oxidative phosphorylation, and negative enrichment in epithelial-mesenchymal transition (EMT) pathways. To reanalyze it with GSPlot, we generated a ranked list by sorting the genes with a descending log-fold-change in their supplementary spreadsheet, and ran enrichment on 10,490 gene sets in the Gene Ontology collection of MSigDB. We obtained much richer results (Fig. 1B,C). Using an FDR cutoff of 0.01, we detected 279 enriched gene sets, organized into 19 clusters. In addition to upregulated oxidative phosphorylation and downregulated EMT, GSPlot identified 4 more upregulated clusters (shown with red labels), and 13 more downregulated clusters (shown with blue labels). Among the upregulated with aging, we find gene set clusters related to muscle contraction, ion transport, mitochondrial ribosome, and ciliary movement, which are completely missed in the original analysis. When we inspect the gene sets in the EMT cluster, we see that one of them is about heart valve development, and the majority of its member genes (43 out of 69) overlap with EMT-related gene sets in the cluster. This suggests downregulation of EMT pathway may be related to shutting down of the developmental pathways in the aging cells.

## 4. Discussion

GSPlot provides a visualization-based approach for interpreting gene set enrichment results. Instead of presenting enriched gene sets as a ranked list, GSPlot organizes them by gene set similarity in a two-dimensional map. This representation makes it easier to inspect groups of biologically related gene sets, especially when the enrichment result contains many overlapping gene sets.

Our reanalysis on aging cardiomyocytes demonstrates the value of running enrichment analysis on high number of gene sets to obtain full resolution, then apply complexity management in visualization, rather than starting with a scaled down collection as in the Hallmark approach. The latter may be preferred for generating very dense visualizations, such as enrichments of multiple cell types in a single table. If more rigorous analysis is needed, however, GSPlot provides a complexity management solution for high-volume results, facilitating easy interpretation and effective communication.

## Acknowledgements

This work is supported by the U.S. National Science Foundation under Grant No. 2440964. TCL was supported by the Clare Boothe Luce Undergraduate Fellowship.

## Notes

### Competing Interest Statement

The authors have declared no competing interest.

